# Individualized cyclic mechanical loading improves callus properties during the remodelling phase of fracture healing in mice as assessed from time-lapsed *in vivo* imaging

**DOI:** 10.1101/2020.09.15.297861

**Authors:** Esther Wehrle, Graeme R Paul, Duncan C Tourolle né Betts, Gisela A Kuhn, Ralph Müller

**Author notes:** **Corresponding authors:** Esther Wehrle, PhD, DVM, Institute for Biomechanics, ETH Zurich, Leopold-Ruzicka-Weg 4, 8093 Zurich, Switzerland, Ralph Müller, PhD, Institute for Biomechanics, ETH Zurich, Leopold-Ruzicka-Weg 4, 8093 Zurich, Switzerland.

## Abstract

Fracture healing is regulated by mechanical loading. Understanding the underlying mechanisms during the different healing phases is required for targeted mechanical intervention therapies. Here, the influence of individualized cyclic mechanical loading on the remodelling phase of fracture healing was assessed in a mouse femur defect model. After bridging of the defect, a loading group (n=10) received individualized cyclic mechanical loading (8-16 N, 10 Hz, 5 min, 3x/week) based on computed strain distribution in the callus using animal-specific real-time micro-finite element analysis. Controls (n=10) received 0 N treatment at the same post-operative time-points. By registration of consecutive scans, structural and dynamic callus morphometric parameters were followed in three callus sub-volumes and the adjacent cortex showing that the remodelling phase of fracture healing is highly responsive to cyclic mechanical loading with changes in dynamic parameters leading to significantly larger callus formation and mineralization. Loading-mediated maintenance of callus remodelling was associated with distinct effects on Wnt-signalling-associated molecular targets Sclerostin and RANKL in callus sub-regions and the adjacent cortex. Given these distinct local protein expression patterns induced by cyclic mechanical loading, the femur defect loading model with individualized load application seems suitable to understand the local spatio-temporal mechano-molecular regulation of the different fracture healing phases.

## Introduction

Mechanical loading is a key factor for normal progression of the fracture healing process. Delayed fracture repair and non-union formation is a major issue in orthopaedic surgery, with an incidence of 5-10% and a high socioeconomic burden ^1,2^ and has been partially attributed to low and inadequate mechanical loading of the defect region ^3,4^, affecting all consecutive phases (inflammation, repair, remodelling) of the fracture repair process. The local mechanical conditions in the fracture healing area (for details see tissue differentiation hypothesis by Claes & Heigele ^5^) are crucial for the repair process by determining molecular and cellular reactions, the tissue to be formed and the type of ossification. We and others have previously shown that externally applied mechanical stimuli, either applied locally to the fracture site via dynamic fixators ^6^ or globally via whole body vibration platforms ^7,8^, can improve fracture healing. In contrast, inadequate mechanical stimuli resulting in a too low or too high interfragmentary motion were not successful or even impaired the healing process ^9–11^; for review see ^12^). In order to understand the underlying mechanisms, refined and well controlled *in vivo* loading models are needed. Recent studies particularly indicate that profound understanding of the local spatio-temporal mechanical regulation of the fracture healing process seems crucial for translation of this knowledge to the clinics, aiming at the development of safe, targeted and individualized mechanical intervention therapies. Whereas many preclinical fracture healing studies focused on improving the early healing phases (inflammation, repair) with the aim of increasing bone formation and achieving earlier bone union, recent studies also indicate a potential to improve long-bone fracture healing outcome via modulation of callus remodelling ^13–15^. Given the strong influence of mechanical loading on normal bone remodelling, the application of controlled mechanical loading might be suitable to also improve and accelerate the remodelling phase during fracture healing. However, preclinical fracture healing studies with application of loading have so far shown contradictory results ^12^ and have not been able to completely capture underlying mechanisms due to restrictions in study design: (I) cross-sectional setup with only end-point analyses, (II) effect assessment not specific to healing phases, (III) same load for all animals irrespective of individual callus properties and healing progression. We aim to overcome current limitations by combining state-of-the-art time-lapsed *in vivo* imaging with a novel precisely controlled femur defect loading system in mice. By applying time-lapsed *in vivo* micro-CT and animal-specific real-time micro-finite element (RTFE) analysis, the developed femur defect loading model allows scaling of loading settings based on strain distribution in the callus^16^, thereby considering individual differences in healing speed. To follow the healing progression in each animal, a recently developed longitudinal *in vivo* micro-CT based approach was applied, which was previously shown to capture the different healing phases and to discriminate physiological and impaired healing patterns^17^, without significant radiation-associated effects on callus properties ^18^. By registering consecutive scans of the defect region for each animal and implementing a two-threshold approach, data on bone turnover and mineralization kinetics can be obtained, which is particularly important for targeting callus remodelling under different conditions.

The objective of this study was to assess the influence of individualized cyclic mechanical loading on the remodelling phase of fracture healing via longitudinal *in vivo* micro-CT imaging. We hypothesized, that individualized cyclic mechanical loading improves structural and dynamic callus properties during the remodelling phase of fracture healing.

The study showed that the remodelling phase of fracture healing is highly responsive to cyclic mechanical loading leading to significantly larger callus formation and mineralization. Loading-mediated maintenance of callus remodelling was associated with distinct effects on Wnt-signalling-associated molecular targets Sclerostin and RANKL in callus sub-regions and the adjacent cortex. The tightly controlled femur defect loading model could widen our knowledge on the local spatio-temporal mechano-molecular regulation of fracture healing relevant for application of mechanical intervention therapies to clinical settings for improvement of impaired healing conditions.

## Results

By combining a novel *in vivo* femur defect loading model (Fig. 1) with a recently established time-lapsed *in vivo* micro-CT based monitoring approach ^18,19^, the influence of individualized cyclic mechanical loading on callus properties was assessed during the remodelling phase of fracture healing. A loading group (n=10) received individualized cyclic mechanical loading (8-16 N, 10 Hz, 5 min, 3x/week) based on computed strain distribution in the callus using animal-specific RTFE, whereas controls (n=10) received 0N for 5 min at the same post-operative time-points. By registration of consecutive scans, structural and dynamic callus parameters were followed in three callus sub-volumes (defect centre: DC, defect periphery: DP, cortical fragment periphery: FP, and the adjacent cortical fragments: FC over time (Fig. 2 and 3). For details on methods see Tourolle et al. 2019 ^19^; for detailed study design see Supplementary Table S1.

**Figure 1.**
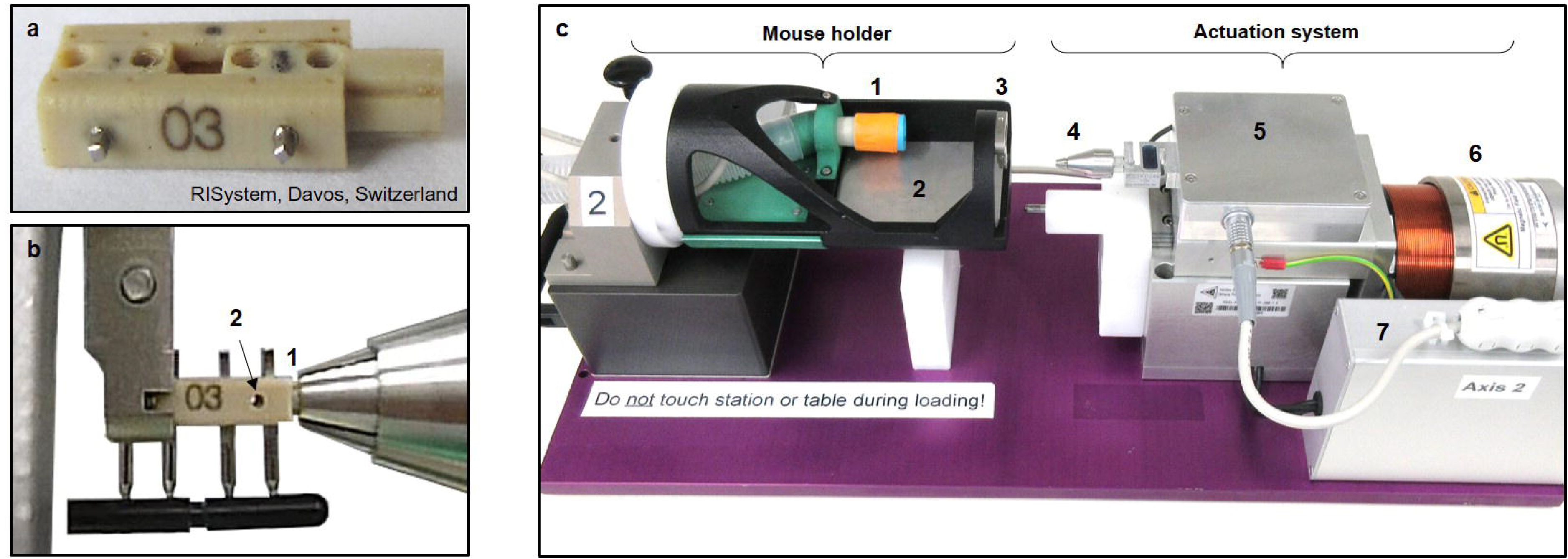
Loading system for femur defects in mice. a. Loading fixator (RISystem, Davos, Switzerland) consisting of four PEEK parts connected via two transverse Titanium pins. b. Insertion of loading fixator into loading device with clamping of loading adapter (1) and removal of transverse pin (2). c. Loading device consisting of mouse holder with anaesthesia inlet (1), heated ground plate (2), clamp for fixator (3) and actuation system with loading chuck (4), load cell electronics box (5), electro-magnetic actuator (6), actuator control electronic box (7). The mouse holder can be used for the loading device and the CT scanner without the need to change the position of the mouse. For detailed description and drawings see Paul et al. ^45^.

**Figure 2.**
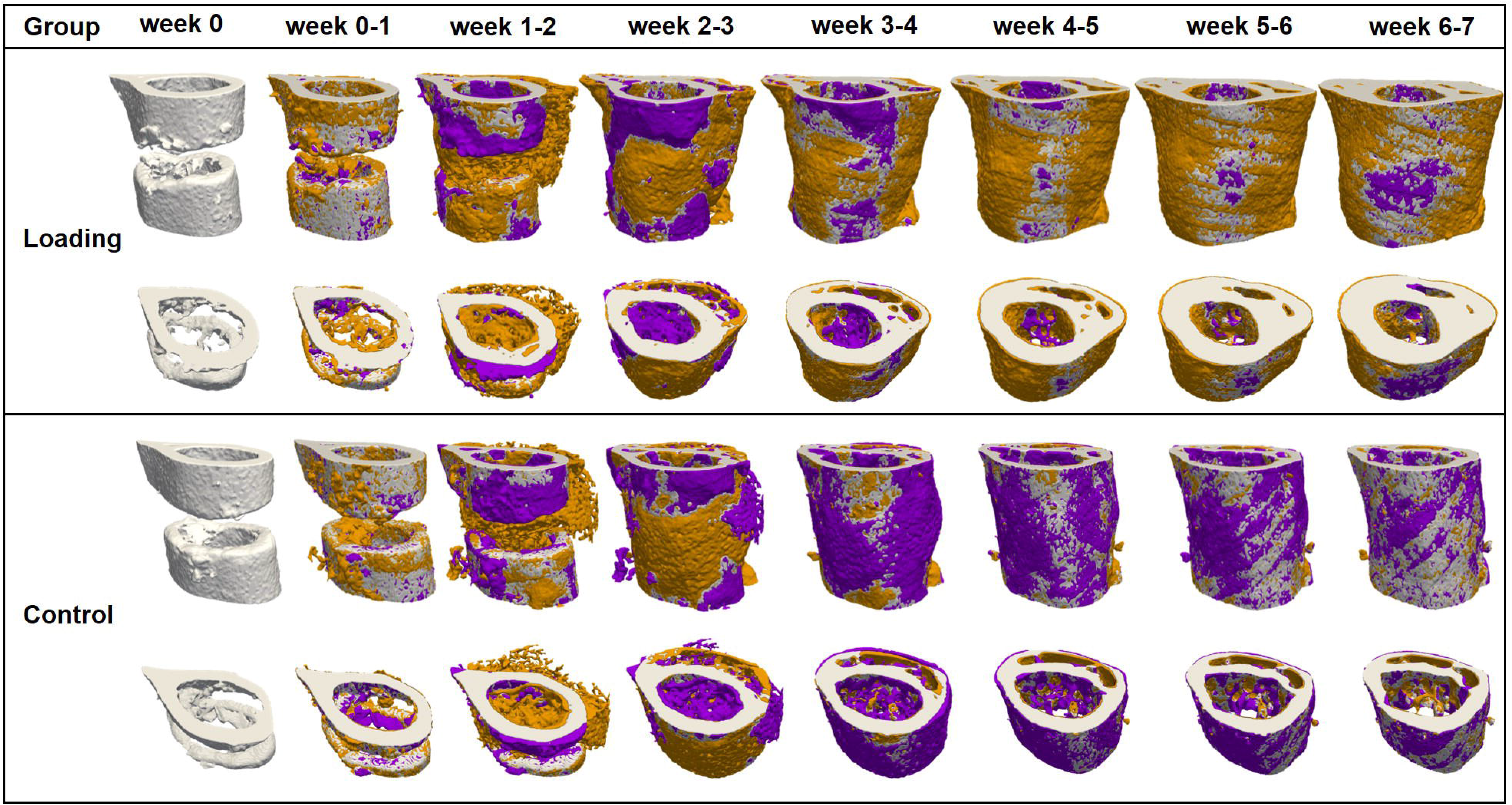
Longitudinal imaging of the defect region via micro-CT. Representative images (full image and cut; threshold: 645 mg HA/cm^3^) of the defect region from animals of the loading group (week 0-7) and the control group (week 0-7). Visualization of bone formation (orange) and resorption (blue) via registration of micro-CT scans from weeks 1-6 to weeks 0-5.

**Figure 3.**
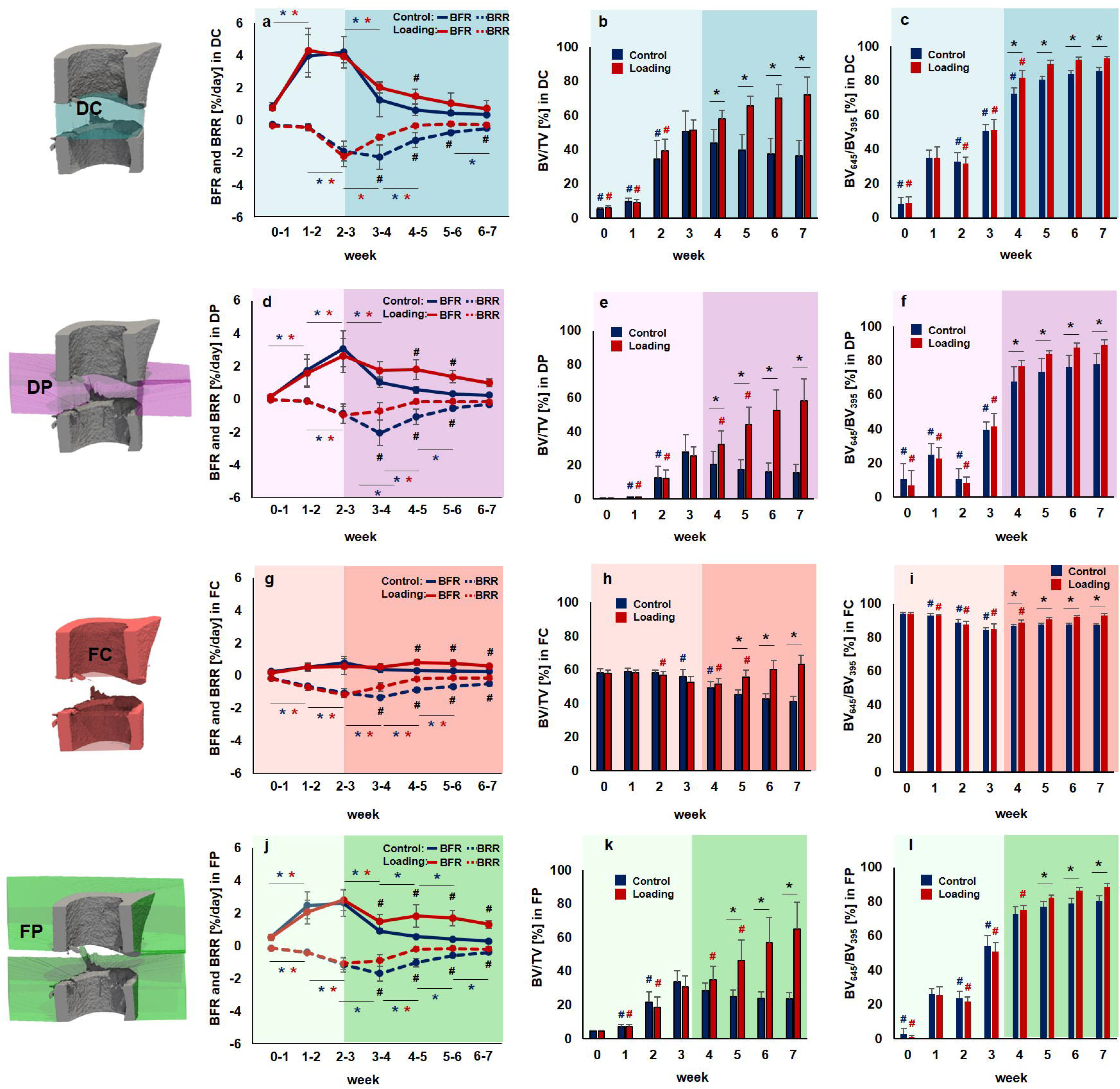
Micro-CT based evaluation of bone parameters in the defect region during the healing period. Four volumes of interest (VOIs) were assessed in the loading group (red) and the control group (blue) from week 0 – week 7 (pre-loading healing period highlighted in lighter colour; loading period highlighted in darker colour): defect center (DC; a-c), defect periphery (DP; d-f), cortical fragment center (FC; g-i), cortical fragment periphery (FP; j-l). a, d, g, j: Bone formation rate (solid line) and bone resorption rate (dashed line) given in percent per day. b, e, h, k: Bone volume (BV) normalized to TV (DC for DC and DP, FC for FC+FP). c, f, i, l: Degree of bone mineralization given as ratio of bone volume with a density ≥645 mg HA/cm^3^ to the total osseous volume (threshold ≥395 mg HA/cm^3^). n=10 per group; * indicates p < 0.05 determined by two-way ANOVA with Geisser-Greenhouse correction and Bonferroni correction.

### General physical observation

All mice recovered rapidly from surgery. During the healing period, the animals’ body weight did not significantly change compared to pre-operative values without significant differences between or within the two groups (see Supplementary Fig. S2). We also assessed the social interaction between mice (animals sitting in groups vs. separated from others) and the nesting behaviour (start of nest building after surgery). Specifically, all animals started to build nests on the day of surgery and were found in groups in the nest in the morning of post-operative day 1. Throughout the further healing period social interaction between mice and nesting behaviour did not differ from pre-surgical observations and was similar for animals of the loading and control groups.

### Volumes of interest (VOI) for evaluation by time-lapsed *in vivo* micro-CT

In order to exclude bias in the further micro-CT analyses, we compared the size of the different VOIs (depicted in Fig. 3; for details on VOI generation see Tourolle et al., 2020 ^19^). The VOIs encompassed the following volume for the control (n=10) and the loading group (n=10): 1.14±0.13mm^3^ vs. 1.04±0.16mm^3^ for the defect centre (DC), 6.68±0.77mm^3^ vs. 6.35±0.82mm^3^ for the defect periphery (DP), 2.28±0.17mm^3^ vs. 2.31±0.16mm^3^ for the cortical fragments (FC), 15.44±0.70mm^3^ vs. 15.84±0.99mm^3^ for the fragment periphery (FP). No significant differences in volume were detected in any of the VOIs between groups.

### Pre-loading healing patterns monitored by time-lapsed *in vivo* micro-CT

In the pre-loading healing period (week 0 to week 3) similar and physiological healing patterns were observed for the control group (n=10) and the loading group (n=10) in all defect VOIs (DC, DP and FP) and the adjacent cortex (FC; Fig. 2 and 3). Registration of consecutive *in vivo* micro-CT scans allowed to capture distinct callus characteristics indicative of the different healing phases (inflammation, repair, remodelling; significant weekly differences in bone parameters indicated in Fig. 3). From week 0-1 to week 1-2 in both groups a significant 4x to 14x increase in bone formation was detected in the three callus VOIs DC (control group: p=0.0007; loading group: p=0.0006; Fig. 3a), DP (control group: p<0.0001; loading group: p<0.0001; Fig. 3d) and FP (control group: p=0.0012; loading group: p=0.0006; Fig. 3j), indicating progression from the inflammation to the reparative phase. This led to a significant gain in bone volume (BV/TV) by week 2 in the callus VOIs DC (control group: p<0.0001; loading group: p<0.0001; Fig. 3b), DP (control group: p<0.0001; loading group: p<0.0001; Fig. 3e) and FP (control group: p<0.0001; loading group: p<0.0001; Fig. 3k). From week 1-2 to week 2-3 a significant 3x to 9x increase in resorptive activities was seen in the callus VOIs DC (control group: p=0.0023; loading group: p<0.0001; Fig. 3a), DP (control group: p=0.0499; loading group: p=0.0162; Fig. 3d) and FP (control group: p=0.0326; loading group: p=0.0070; Fig. 3j), indicating the progression from the repair to the remodelling phase. From week 2 to week 3, the highly mineralized bone fraction in the callus VOIs significantly increased (p<0.0001 in DC, DP, FP) in both groups, indicating callus maturation (Fig. 3c, 3f and 3l).

From week 0 to week 3, in both groups, strong resorptive activities (Fig. 2 and 3g) were seen in the adjacent cortical fragments (FC). BRR showed significant weekly increases whereas no significant weekly changes in bone formation were observed in this VOI. The strong resorptive activities let to decreased bone volume by week 3 reaching statistical significance in the loading group (p<0.0001). Furthermore, the fraction of highly mineralized bone significantly decreased from week 1 to week 3 in both groups (p<0.0001), indicating cortical reorganisation.

### Individualized cyclic mechanical loading improves callus properties during the remodelling phase of fracture healing

Individualized cyclic mechanical loading (8-16N, 10 Hz, 300 cycles, 3x/week) started after bridging of the femur defect (week 3; Table 1) at the beginning of the remodelling phase (loading group, n=10). RTFE allowed determination of initial loads (8-12N) based on individual callus properties, weekly load-scaling (8-16N) and matching of strain distribution in the callus for the 4-week loading period. Control animals received 0N loading at the same post-operative time-points. Already after 1 week, the first loading-associated effects on callus properties and the adjacent cortical bone were seen: Specifically, loading reduced BRR in all VOIs in week 3-4 compared to pre-loading values in week 2-3 (Fig. 3a, d, g, j) reaching statistical significance in DC (p<0.0001). In contrast, in the control group, BRR further increased in all VOIS during the same period with statistical significance seen in the peripheral VOIs (p=0.0066 for DP, p=0.0126 for FP). From week 3-4 to week 6-7, the applied individualized mechanical loading was also able to maintain bone formation at significantly higher levels compared to controls, while bone resorption was significantly reduced during the same period. This led to a linear increase in osseous callus volume in loaded animals from week 3 to week 7 in all callus VOIs (DC: +41 %, DP: +130 %, FP: +111 %) and the adjacent cortex (FC: +20 %), whereas it significantly declined in controls (DC: - 28 %, DP: - 44 %, FP: −27 %, FC: −30 %). In week 6-7, no significant differences in bone turnover were seen anymore indicating load adaptation. After 7 weeks, loaded animals had significantly more bone in the defect (2x for DC, p<0.0001; 4x for DP, p<0.0001; 3x for FP, p<0.0001) with a significantly larger fraction of highly mineralized bone compared to controls (+8% for DC, p=0.0006; +15% for DP, p<0.0015; +10% for FP, p<0.0001). Three-point bending also showed a 46% higher relative bending stiffness (healed femur vs. contralateral femur) in the loaded animals (107%) compared to controls (73%; n=2/group; Supplementary Fig. S3). In addition, according to the standard clinical evaluation of X-rays, the number of bridged cortices per callus was evaluated in two perpendicular planes and animals with ≥3 bridged cortices were categorized as healed. From week 4 to week 7 all defects in both groups were categorized as healed (Table 1), indicating that the applied forces during cyclic mechanical loading did not lead to re-fracture of the callus.

**Table 1.**
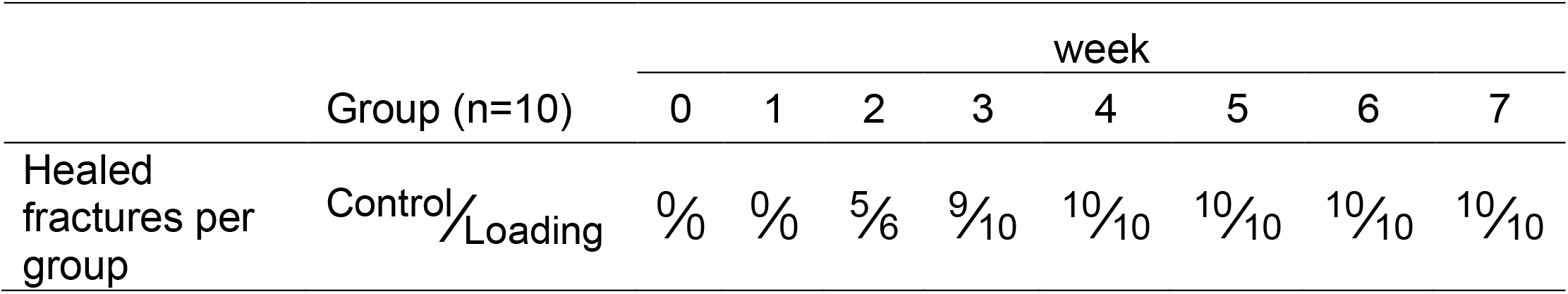
Fracture healing outcome assessed by cortical bridging (healed fracture ≥ 3 bridged cortices, threshold 395 mg HA/cm^3^).

Loading also affected the bone turnover in the adjacent cortex (FC). Already after 1 week, loaded animals showed significantly lower cortical bone resorption compared to controls (Fig. 3 j). Whereas cortical volume significantly declined (p<0.0001) in control animals from week 3 to week 7 by 27%, it significantly increased (p<0.0001) by 20% in the loaded animals during the same time period (Fig. 3k). From week 4 to week 7 loading was also associated with significantly higher cortical mineralization (+4% in week 4 to +54% in week 7) compared to controls (week 4: p=0.0459, week 5 – week 7: p<0.0001; Fig. 3l).

In week 7, the FC VOI comprised 46% and 34% of the osseous tissue in the total VOI (TOT) for the control and loading group, respectively. In the DC VOI, 20% (control group) and 17% (loading group) of the total bone volume were seen. In the two peripheral VOIs, 9% (control group) and 14% (loading group) of the total osseous tissue were seen in the DP and 26% (control group) and 35% (loading group) in the FP VOI. Loading was associated with a shift in the bone distribution from the cortical fragments to the peripheral VOIs.

### Histology

In order to also visualize the tissue composition of the callus, we performed end point histological stainings of serial sections from one animal of each group. Histology supported our micro-CT findings with complete cortical bridging seen in the animal from the loading and from the control group (Fig. 4 B-I). However, the loaded animal showed a larger callus with some cartilage fractions suggesting ongoing endochondral ossification processes, whereas in the control animal the callus was largely remodelled with restoration of the medullary cavity indicating proceeding towards the end of healing (Fig. 4 b-e). Furthermore, differences in the shape of the cell nuclei were observed between the osteocytes in the fracture callus (round shape) of the loaded animal in comparison to the cell nuclei seen in the cortical bone of the same animal (ellipsoidal shape) and the cell morphology seen in the control animals (Fig. 5 a+b), suggesting an association between the local mechanical environment and cellular morphology. In regions with lower strains visualized by micro-FE (endosteal fracture callus; Fig. 4a and 5b), rounder cell nuclei were observed, compared to ellipsoidal nuclei shape in higher strained regions (cortex; Fig. 4a and 5b). To capture potential underlying mechano-molecular targets, we performed immunohistochemistry of Sclerostin (inhibitor of the mechano-responsive and osteoanabolic Wnt signalling pathway) and RANKL (negatively-regulated target gene of Wnt signalling) which have both previously been associated with the mechanical regulation of bone adaptation and healing ^8,20–24^. Less abundant Sclerostin staining was visible in the fracture callus compared to the cortical bone of the loaded animal (Fig. 4 f and 5b) with no region-specific differences in staining patterns being seen in the control animal (Fig. 4g and Fig. 5b), suggesting that Wnt-signalling contributes to the loading-mediated osteoanabolic effects on fracture healing seen in this study. At the same time RANKL, which is down-regulated by Wnt-signalling, was more abundantly and stronger expressed in the callus of the loaded animal compared to the control animal (Fig. 4 h+i and Fig. 5b). In line with previous studies, strong RANKL expression was seen in lowly strained callus regions as visualized via micro-FE analysis (Fig. 4a and 5b), indicating ongoing bone remodelling.

**Figure 4.**
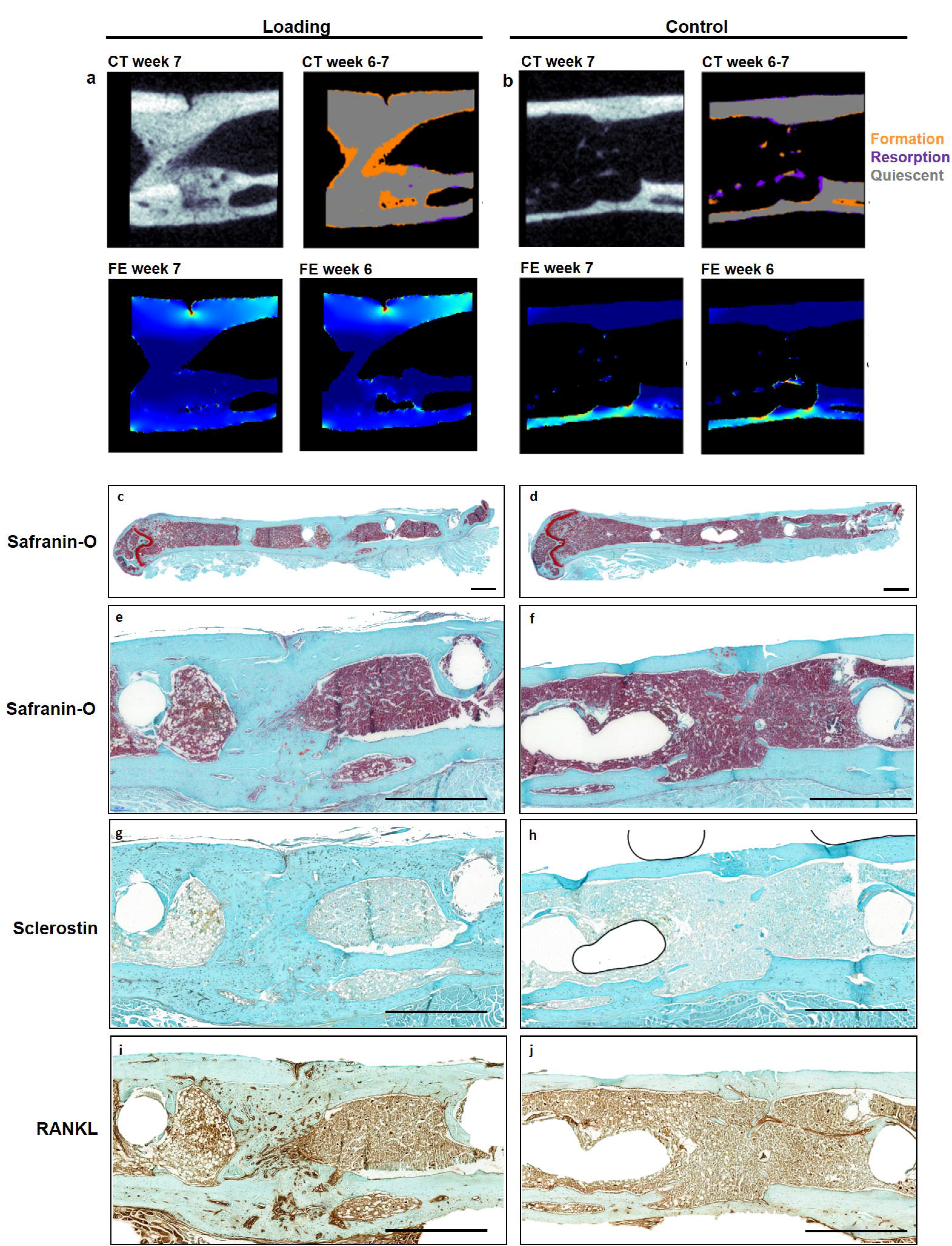
Micro-CT and histological analyses of consecutive callus sections. a, b: Micro-CT images of fracture callus sections corresponding to sections stained with Safranin-O for one animal from the loading (a) and the control group (b): CT image (threshold: 395 mg HA/cm^3^), visualization of regions of bone formation and resorption via registration of CT image from weeks 6 and 7, visualization of strains on CT section via micro-FE analysis for week 7 and week 6. c-j. Consecutive longitudinal sections of fractured femora 7 weeks after defect surgery of one animal from the loading group and the control group stained for Safranin-O (c-f), Sclerostin (g, h), RANKL (i, j). Scale bar = 100 μm (c, d), scale bar = 500 μm (e-j).

**Figure 5.**
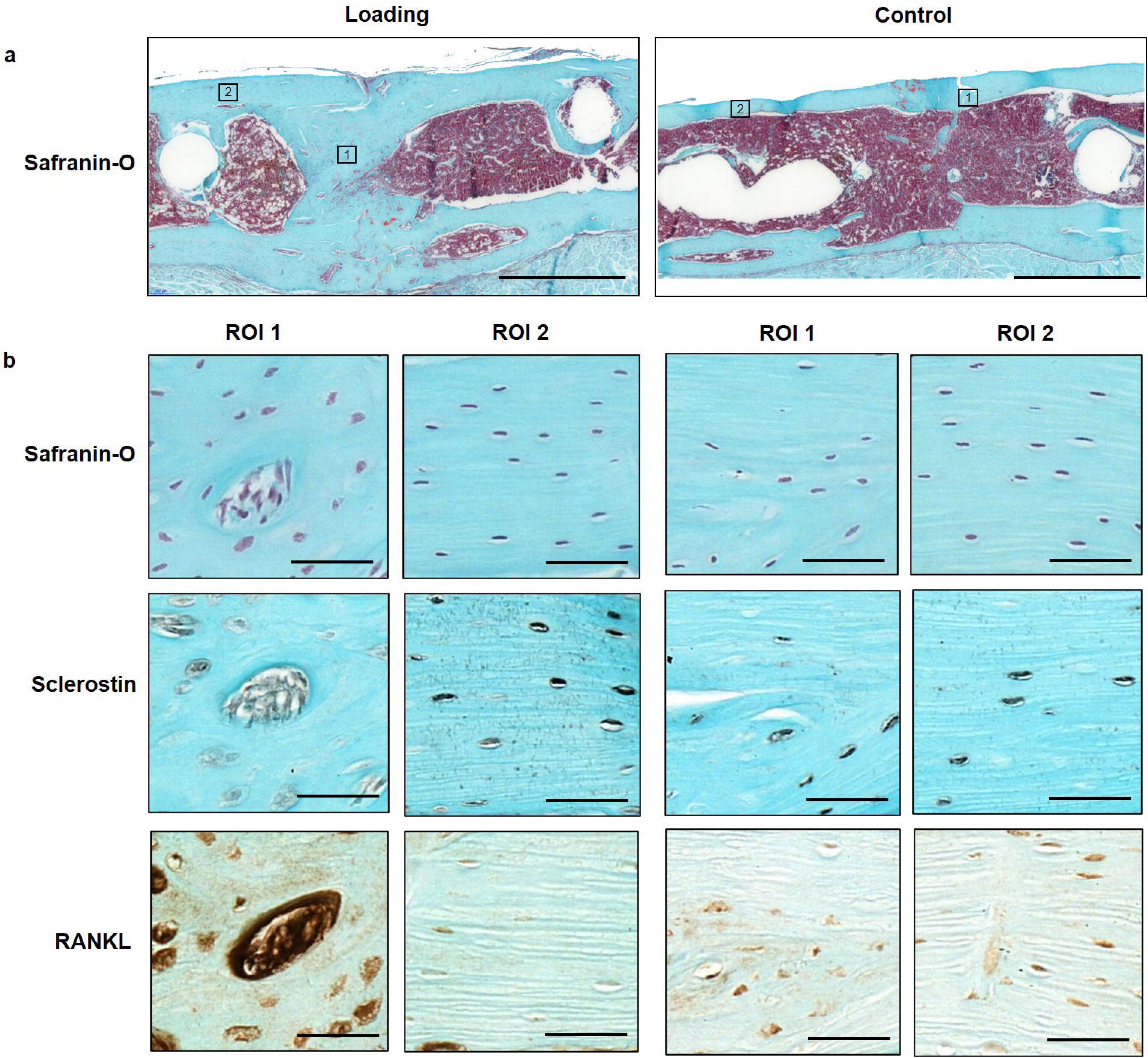
Visualization of local cellular distribution patterns and protein expression in the fracture callus and adjacent cortex. a. Longitudinal Safranin-O stained sections of fractured femora 7 weeks after defect surgery of one animal from the loading group and one animal from the control group. Boxes indicate regions depicted in b (1: callus; 2: cortex); Scale bar = 100 μm. b. Magnification of callus region (left) and cortical region (right) stained for Safranin-O, Sclerostin and RANKL; Scale bar = 50 μm.

## Discussion

In this study, the effect of individualized cyclic mechanical loading on the remodelling phase of fracture healing was assessed in a novel femur defect loading model in mice using a recently developed *in vivo* micro-CT based monitoring approach. Given the strong mechano-responsiveness of bone cells, several studies have applied mechanical loading (e.g. globally via vibration platforms, or locally via external fixators) to improve fracture healing showing contradictory results. Whereas some studies saw a loading-mediated improvement of fracture healing, others were not successful or even reported an impairment of the fracture healing process ^9–11^. One limitation of these studies was the application of the same load to all animals irrespective of individual differences in healing progression, which bears the risk of re-fracturing lowly mineralized bone after initial bridging of the defect. In order to allow for individualized mechanical loading we now developed a precisely controlled *in vivo* loading model based on external fixation of femur defects in mice. By combining state-of-the-art time-lapsed *in vivo* imaging with individualized mechanical loading protocols adapted to animal-specific callus properties, this allows to precisely characterize spatio-temporal effects of mechanical loading on the fracture callus. Given the high mechano-responsiveness of bone cells during normal bone remodelling and as recent studies also indicate a potential to improve fracture healing outcome via modulation of callus remodelling ^13–15^, we focused this study on the effect of mechanical loading on callus properties during the remodelling phase of fracture healing.

Via weekly *in vivo* micro-CT imaging, we saw, that cyclic mechanical loading linearly increased the osseous callus volume during the remodelling phase of fracture healing, so that by the study end, loaded animals had significantly more bone in all callus VOIs (2x for DC, p<0.0001; 4x for DP, p<0.0001; 3x for FP, p<0.0001) with a significantly larger fraction of highly mineralized bone compared to controls (+8% for DC, p=0.0006; +15% for DP, p<0.0015; +10% for FP, p<0.0001). This is supported by recent study from Liu et al. ^25^ describing additional loading-mediated bone formation during the remodelling phase in a uni-cortical tibia defect model in mice. However, they failed to show statistical significance between the loading and the control group in histomorphometry and micro-CT analyses, which they attributed to the high variability within groups. As the study applied the same load to all animals irrespective of healing progression, differences in the induced strains might have caused the high variability within groups. We addressed this by using our recently developed RTFE based loading approach ^26^, which takes into account the individual strain distribution within the callus for determining individual loading settings at weekly post-operative intervals. In a recent study by Malhotra et al. ^27^ individualized cyclic mechanical loading (week 0 – week 4, 3x/week) was shown to significantly increase bone volume fraction in a partial vertebral defect model compared to controls. So far, no study using a complete defect model assessed the effect of local cyclic mechanical loading (e.g. via external fixators, tibia/ulna-loading) when exclusively applied during the remodelling phase of the healing process. An earlier tibia defect loading study (defect size: 3 mm) in sheep by Goodship et al. ^6^ applied cyclic mechanical loading (25 μm interfragmentary movement, 30 Hz) throughout the healing period (week 0 - week 10, 5x per week) and reported a significantly larger callus with higher mineralization in loaded animals compared to controls. However, as the loading application was not restricted to specific post-operative time frames, it was not possible to evaluate whether loading during a specific fracture healing period was sufficient for the observed improvement in fracture healing or if cyclic mechanical loading needs to be applied throughout the healing process to achieve such an effect. We now showed, that the remodelling phase is highly responsive to cyclic mechanical loading applied after initial cortical bridging of the defect, with pronounced effects on callus size and mineralization. Via limiting the loading application to the earlier healing period (day 7 – day 19), Gupta et al. ^28^ aimed at assessing (histomorphometry and radiography on day 42) whether there is a sustained effect of cyclic mechanical loading (2%, 10% and 30% strain, 0.5 Hz, 500 cycles) on fracture healing outcome. They found a larger bone fraction in the callus and higher cortical bridging scores in the loaded compared to the control animals, indicating, that loading effects persisted after a 4-week non-loading period. In contrast to these studies, Schwarz et al. ^29^ did not observe any loading-mediated improvement on fracture healing in a critically-sized femur defect model in rats (defect size: 5mm). However, in this study loading (6 cycles of constant loading at 500 μm IFM and 40s rest phase), was only applied once per week, which indicates a critical role of the type of loading (constant vs. cyclic) and the frequency of loading sessions per week. Whether cyclic mechanical loading in contrast to constant mechanical loading is able to prevent non-union formation in critically-sized defects needs to be assessed in future studies.

We were now able to also include dynamic parameters such as bone formation and resorption in the monitoring of loading-mediated effects on fracture healing via registration of longitudinal *in vivo* micro-CT scans (for details see Tourolle et al. 2019 ^19^). Compared to histology-based determination of the bone formation rate via *in vivo* application of fluorescent dyes (e.g. calcein, alizarin), our micro-CT-based monitoring approach with registration of consecutive scans also allows for assessment of the bone resorption rate in individual animals, which was shown to be particularly suited for the characterization of the remodelling phase during the fracture healing process ^18,30^. The most striking loading-mediated effect in the current study was the prompt and significant reduction in bone resorption rate (week 3-4 vs. week 2-3), which was seen in the central and periosteal VOIs (DC, FP) as early as 1 week after loading (Fig. 3A, 3J). Compared to control animals, in which bone resorption still increased during this early remodelling period (week 3-4), loaded animals showed significantly lower bone resorption rates in all callus VOIs (DC, DP, FP) in this first week of loading, indicating a strong mechano-responsiveness of the fracture callus during this healing period. Most of the previous studies either assessed the loading effects only after a longer loading period of minimum 2 weeks ^29,31^ or they did not see significant loading-mediated effects on the fracture callus after shorter treatment periods ^8,10,28^. So far only one study by Leung et al. ^32^ was able to detect a loading-mediated improvement of fracture healing (larger callus diameter) after a short loading period of 1 week. We were now able to strengthen these cross-sectional structural callus findings with our longitudinal micro-CT data obtained for single animals. In respect to the underlying mechanisms, we showed for the first time, that not only bone formation but also bone resorption can be significantly modulated via cyclic mechanical loading during fracture healing, indicating a potential for mechanical intervention therapies to improve impaired fracture healing conditions associated with imbalanced bone remodelling. Furthermore, implementation of our two-threshold approach showed cyclic mechanical loading to be also effective in significantly advancing callus mineralization compared to control animals. Via combining our novel loading model with time-lapsed *in vivo* imaging and a two-threshold approach, we were able to characterize the spatio-temporal effects of loading during the remodelling phase of fracture healing. Furthermore, the applied micro-FE-based loading approach also allowed for spatio-temporal visualization of strains induced in the fracture callus (Fig. 4a+b). In order to also visualize the tissue composition of the callus, we performed end point histological stainings of serial sections from one animal of each group. Histology supported our micro-CT findings with complete cortical bridging seen in both groups. However, the loaded animal showed a larger callus with some cartilage fractions suggesting ongoing endochondral ossification processes, whereas in the control animal the callus was largely remodelled with restoration of the medullary cavity indicating proceeding towards the end of healing. Furthermore, differences in the shape of the cell nuclei were seen between the osteocytes in the fracture callus (round shape) of the loaded animal in comparison to the cell nuclei seen in the cortical bone of the same animal (ellipsoidal shape) and the cell morphology seen in the control animals. This is particularly interesting, as previous *in vitro* studies found, that round cells are more sensitive to mechanical loading compared to flat and elongated cells ^33–35^, suggesting that repeated cyclic mechanical loading might be able to induce round cellular morphology of (early) osteocytes, thereby maintaining the remodelling phase of fracture healing eventually leading to higher mineralization of the newly formed tissue compared to controls. In the *in vivo* setting osteocytes reside in lacunae within the lacunocanalicular network (LCN). A recent study by Casanova et al. ^36^ longitudinally assessed the development and evolution of the LCN during fracture healing indicating a progressive increase in the complexity of the LCN, which they attributed to factors expressed by osteocytes such as matrix metalloproteinases, Sclerostin and RANKL. To capture potential underlying molecular targets of the spatio-temporal loading effects seen in the current study, we performed immunohistochemistry of Sclerostin (inhibitor of the mechano-responsive and osteoanabolic Wnt signalling pathway) and RANKL (negatively-regulated target gene of Wnt signalling) which have both previously been associated with the mechanical regulation of bone adaptation and healing ^8,20–24^. Whereas visual inspection showed less abundant Sclerostin staining in the fracture callus compared to the cortical bone of the loaded animal (Fig. 4f, Fig. 5b), no region-specific differences in staining patterns were seen in the control animal (Fig. 4g, Fig. 5b). This might indicate that Wnt-signalling contributes to the loading-mediated osteoanabolic effects on fracture healing seen in this study. However, at the same time we saw more and stronger RANKL staining in the callus of the loaded animal compared to the control animal (Fig. 4 h+i). This finding precludes that the loading-mediated increases in RANKL expression seen in this study were solely mediated via Wnt-signalling but rather an interplay with further signalling pathways must have triggered this strong response in RANKL production. Via micro-FE analysis, we saw high RANKL expression in the lowly strained endosteal fracture callus, indicating ongoing bone remodelling. In order to shed light into the complex interplay between a multitude of mechano-responsive molecules and signalling pathway, novel advents in RNAsequencing including spatial transcriptomic approaches might, in future studies, open interesting possibilities to understand the spatio-temporal mechanomics of fracture healing. In non-fractured bone, some of these molecular technologies (e.g. microarrays, ^37–40^) have already been successfully applied to study mechano-molecular effects in well-controlled and characterized loading models (e.g. ulna, tibia and tail loading) in rodents. Targets (e.g. Sclerostin, Estrogen signalling) found via these comprehensive studies were then assessed in detail in further studies using either models with local ^41–43^ or global load application ^44^. As the main underlying mechanisms of bone adaptation to loading were identified using local loading models of long bones (ulna, tibia ..), this indicates that comprehensive spatio-temporal mechano-molecular analyses in fracture healing will require tightly regulated models with local application of mechanical loading being more controllable and less prone to variation induced by confounding factors compared to models with global loading application.

In summary, using our recently developed femur defect loading model in combination with time-lapsed *in vivo* micro-CT and a two-threshold approach, we showed that the remodelling phase of fracture healing is highly responsive to cyclic mechanical loading with changes in dynamic callus parameters leading to larger callus formation and mineralization. Loading-mediated maintenance of callus remodelling was associated with distinct effects on the Wnt-signalling-associated molecular targets Sclerostin and RANKL in callus sub-regions and the adjacent cortex as assessed via immunohistochemistry. Given these distinct local protein expression patterns induced by external cyclic mechanical loading, our tightly-controlled femur defect loading model with individualized load application could be used in future studies to precisely understand the local spatio-temporal mechano-molecular regulation of the different fracture healing phases. With our RTFE based approach for individualized load application considering differences in defect geometry and healing progression, the femur defect loading model could also be used to study the bone regeneration capacity of biomaterials under load application. With further advents in transcriptomics and registration techniques for different imaging technologies, the tightly controlled femur defect loading model could widen our knowledge on the local spatio-temporal mechano-molecular regulation of fracture healing relevant for application of mechanical intervention therapies to clinical settings for improvement of impaired healing conditions.

## Methods

### Study design

By combining a novel *in vivo* femur defect loading model with a recently established time-lapsed *in vivo* micro-CT based monitoring approach, the influence of individualized cyclic mechanical loading on callus properties was assessed during the remodelling phase of fracture healing. All mice received a femur defect and post-operative micro-CT scans (vivaCT 40, ScancoMedical, Brüttisellen, Switzerland) were performed. After bridging of the femur defect (week 3), RTFE allowed determination of initial loads (8N-12N), and weekly load-scaling and matching of strain distribution in the callus for the 4 week loading period. The loading group (n=10) received individualized cyclic mechanical loading (8-16N, 10 Hz, 5 min, 3x/week) whereas controls (n=10) received 0N for 5 min at the same post-operative time-points. By registration of consecutive scans, structural and dynamic callus parameters were followed in three callus sub-volumes (defect centre: DC, defect periphery: DP, cortical fragment periphery: FP) and the adjacent cortical fragments (FC) over time (Figure 3, for details on methods see Tourolle et al. 2020^19^; for detailed study design see Supplementary Table S1).

### Animals

All animal procedures were approved by the Commission on Animal Experimentation (license number: 181/2015; Kantonales Veterinäramt Zürich, Zurich, Switzerland). We confirm that all methods were carried out in accordance with relevant guidelines and regulations (Swiss Animal Welfare Act and Ordinance (TSchG, TSchV)) and reported considering ARRIVE guidelines. To study adult fracture healing female 12 week-old C57BL/6J mice were purchased from Janvier (Saint Berthevin Cedex, France) and housed in the animal facility of the ETH Phenomics Center (EPIC; 12h:12h light-dark cycle, maintenance feed (3437, KLIBA NAFAG, Kaiseraugst, Switzerland), 5 animals/cage) for 8 weeks. At an age of 20 weeks, all animals received a femur defect by performing an osteotomy with a 0.66mm Gigli wire saw as previously described (group 1: control group, n=10; group 2: loading group, n=10; housing after surgery: 2-3 animals/cage; for details on study design see Supplementary Table 1). All defect surgeries were performed by the same veterinarian. Perioperative analgesia (25 mg/L, Tramal^®^, Gruenenthal GmbH, Aachen, Germany) was provided via the drinking water two days before surgery until the third post-operative day. For surgery and micro-CT scans, animals were anaesthetized with isoflurane (induction/maintenance: 5%/1-2% isoflurane/oxygen). Perioperative handling, and monitoring, micro-CT imaging and loading application was performed by the surgeon.

### Loading fixator assembly and group assignment

The four parts of the loading fixators (n=20) were assembled as depicted in Fig. 1. To allow optimal identification throughout the experiments, one side part was engraved with a fixator-specific number (Fig 1a, 1b). The stiffness of each fixator was measured using a Zwick testing machine and the fixators were assigned to the two groups, to allow similar distributions of fixator stiffness in the loading and control group (Supplementary Fig. 3).

### Femur osteotomy

In all animals an external fixator (Mouse ExFix, RISystem, Davos, Switzerland; mean stiffness: 17N/mm; Suppl. Fig. 1) was positioned at the craniolateral aspect of the right femur and attached using four mounting pins. First, the most proximal pin was inserted approximately 2mm proximal to the trochanter, followed by placement of the most distal and the inner pins. Subsequently, a femur defect was created using a Gigli wire (diameter: 0.66mm).

### Time-lapsed *in vivo* micro-CT

Immediate post-surgery correct positioning of the fixator and the defect was visualized using a vivaCT 40 (Scanco Medical AG, Brüttisellen, Switzerland) (isotropic nominal resolution: 10.5 μm; 2 stacks of 211 slices; 55 kVp, 145 μA, 350 ms integration time, 500 projections per 180°, 21 mm field of view (FOV), scan duration ca. 15 min). Subsequently, the fracture callus and the adjacent bone between the inner pins of the fixator were scanned weekly using the same settings. Scans were registered consecutively using a branching scheme (registration of whole scan for bridged defects; separate registration of the two fragments for unbridged defects)^5^. Subsequently, morphometric indices (bone volume - BV, bone volume/total volume – BV/TV, bone formation rate – BFR, bone resorption rate - BRR) were computed (threshold: 395mg HA/cm^3^; for details on methods see Tourolle et al., 2020 ^19^). To assess mineralization progression, a second threshold (645mg HA/cm^3^) was applied and the ratio between highly and lowly mineralized tissue (BV645/BV395) was calculated. The two selected thresholds are included in our recently developed multidensity threshold approach ^19^. According to the standard clinical evaluation of X-rays, the number of bridged cortices per callus was evaluated in two perpendicular planes (UCT Evaluation V6.5-1, Scanco Medical AG, Brüttisellen, Switzerland). A “healed fracture” was considered as having a minimum of at least three bridged cortices per callus.

For evaluation, four volumes of interest (VOIs) were defined, which were created automatically from the post-operative measurement (Fig. 3): defect centre (DC), defect periphery (DP), cortical fragment centre (FC), and fragment periphery (FP). Data were normalised to the central VOIs: DC/DC, DP/DC, FC/FC, FP/FC.

### Individualized cyclic mechanical loading

From week 4 to week 7, individualized cyclic loading (8-16N, 10Hz, 3000cycles; 3x/week; controls - 0N) was applied via the external loading fixator (Fig 1b) based on computed strain distribution in the callus using animal-specific RTFE analysis (for detailed description of methods see ^26^). Briefly, after weekly micro-CT measurements of each animal, the images were pre-processed using threshold-binning to create a high-resolution multi-density FE mesh, which was then solved on a supercomputer within the same anaesthetic session for each mouse. 2D and 3D visualizations of the callus were generated and a strain histogram was plotted. This information allowed load scaling based on the strains induced in the callus in individualized animals. Specifically, the distribution was scaled to achieve a median effective microstrain of 700. To assess regions that posed a failure risk the 99th percentile strains of each simulation was visualised in both image based and histogram forms. If more than 50 voxels exceeded 1% then the optimised load was reduced by 1 N.

### Mechanical testing

3 point bending tests were performed on a Zwick compression tester (ZwickRoell GmbH & Co, Ulm, Germany). Tests were conducted on a setup of 8 mm support span and up to 3 N. Tests were performed quasi-statically with a cross-head speed of 0.4 mm/min. Stiffness was then calculated from the linear region of the resultant force-displacement curve. Each sample was tested three times and the mean stiffness was taken.

### Histology

Histological stainings were performed in one animal per group. On day 49, femora were excised, the femoral head was removed and the samples were placed in 4% neutrally buffered formalin for 24 hours and subsequently decalcified in 12.5% EDTA for 10-14 days. The samples were embedded in paraffin and the complete fracture callus was cut in 10 μm longitudinal serial sections. Every 10th section was stained with Safranin-O: Weigert’s iron haematoxylin solution (HT1079, Sigma-Aldrich, St. Louis, MO) - 4min, 1:10 HCl-acidified 70% ethanol - 10s, tap water - 5min, 0.02% Fast Green (F7258, Sigma-Aldrich, St. Louis, MO) - 3min, 1% acetic acid - 10s, 0.1% Safranin-O (84120, Fluka, St. Louis, MO) - 5min. Images were taken with Slide Scanner Pannoramic 250 (3D Histech, Budapest, Hungary) at 20x magnification. Sections in between were stained for Sclerostin and RANKL.

For immunohistochemical staining of Sclerostin, nonspecific sites were blocked (1% BSA/PBS + 1% rabbit serum) for 60 min at room temperature. Subsequently, the sections were incubated with the primary antibody against Sclerostin (AF1589, R&D Systems, Minneapolis, MN; 1:150 in 1%BSA/PBS + 0.2% rabbit serum) overnight at 4°C. To detect the primary antibody, a secondary biotinylated rabbit anti-goat-IgG antibody (BAF017, R&D Systems, Minneapolis, MN) was added for 1 h at room temperature. For signal amplification, the slides were incubated with avidin-biotin complex (PK-6100 Vector Laboratories, Burlingame, CA) for 30 min. Diaminobenzidine (Metal Enhanced DAB Substrate Kit, 34065 ThermoFisher Scientific, Waltham, MA) was used as detection substrate. Counterstaining was performed with FastGreen (F7258, Sigma-Aldrich, St. Louis, MO).

For immunohistochemical staining of RANKL, slides were placed into TBS-Tween buffer and inserted into the Dako-Autostainer (Dako, Ft. Collins, USA) using the following staining protocol: incubation with primary antibody against RANKL (1:100; ab 9957, Abcam, Cambridge, UK) at 4°C overnight, rinsing with TBS-Tween, peroxidase blocking for 10min at room temperature, rinsing with TBS-Tween, Envision+System HPR Rabbit (Dako K4003) for 60 min at room temperature, rinsing with TBS-Tween and incubation with DAB (Dako K3468) for 10 min at room temperature. The slides were rinsed in A. dest and counterstained with FastGreen for 2sec.

For both immunohistochemical stainings, species-specific IgG was used as isotype control. Images were taken with Slide Scanner Pannoramic 250 (3D Histech, Budapest, Hungary) at 40x magnification.

### Statistics

Data were tested for normal distribution (Shapiro-Wilk-Test) and homogeneity of variance (Levene-Test). Depending on the test outcome, group comparisons (loading vs. control group) of data derived at single time points (VOI size) were performed by two-tailed Student’s t-test or Mann-Whitney U-test (IBM SPSS Statistics Version 23). For statistical evaluation of repeated measurements two-way ANOVA with Geisser-Greenhouse correction and Bonferroni correction (GraphPad Prism 8) were performed. The level of significance was set at p < 0.05.

## Acknowledgements

We thank Marco Hitz and Peter Schwilch for technical assistance. We thank Dr. Angad Malhotra and Jianhua Zhang for surgery assistance. We thank Christopher M. O’Neill for the micro-CT and micro-FE images depicted in Figure 4. The authors gratefully acknowledge support from the EU (ERC Advanced MechAGE ERC-2016-ADG-741883). E. Wehrle received funding from the ETH Postdoctoral Fellowship Program (MSCA-COFUND, FEL-25_15-1).

## Author Contributions Statement

The study was designed by E.W., G.A.K., and R.M.. The experiments were performed by E.W. and G.R.P. Data analyses were performed by E.W., G.R.P. and D.C.B.. The manuscript was written by E.W. and reviewed and approved by all authors.

## Data availability

All necessary data generated or analysed during the present study are included in this published article and its Supplementary Information files (preprint available on BioRxiv (BIORXIV/2020.09.15.297861). Additional data that support the findings of this study are available from the corresponding author upon reasonable request.

