## Supplementary Information for "Individualized cyclic mechanical loading improves callus properties during the remodelling phase of fracture healing in mice as assessed from time-lapsed *in vivo* imaging"

**Corresponding authors:**

Esther Wehrle, PhD, DVM

8093 Zurich, Switzerland

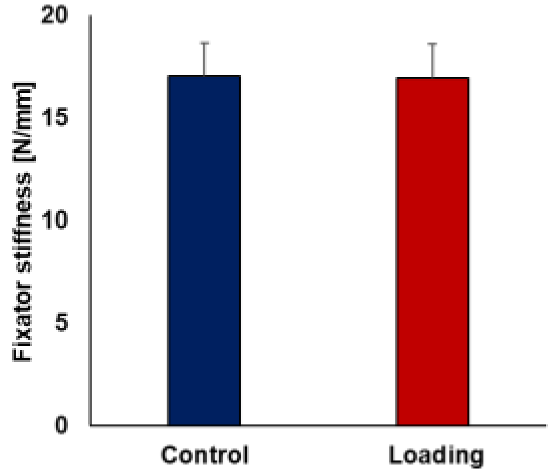


**Supplementary Fig. S1.** Pre-operative measurements of fixator stiffness were used to assign fixators to the loading group (n=10) and the control group (n=10).

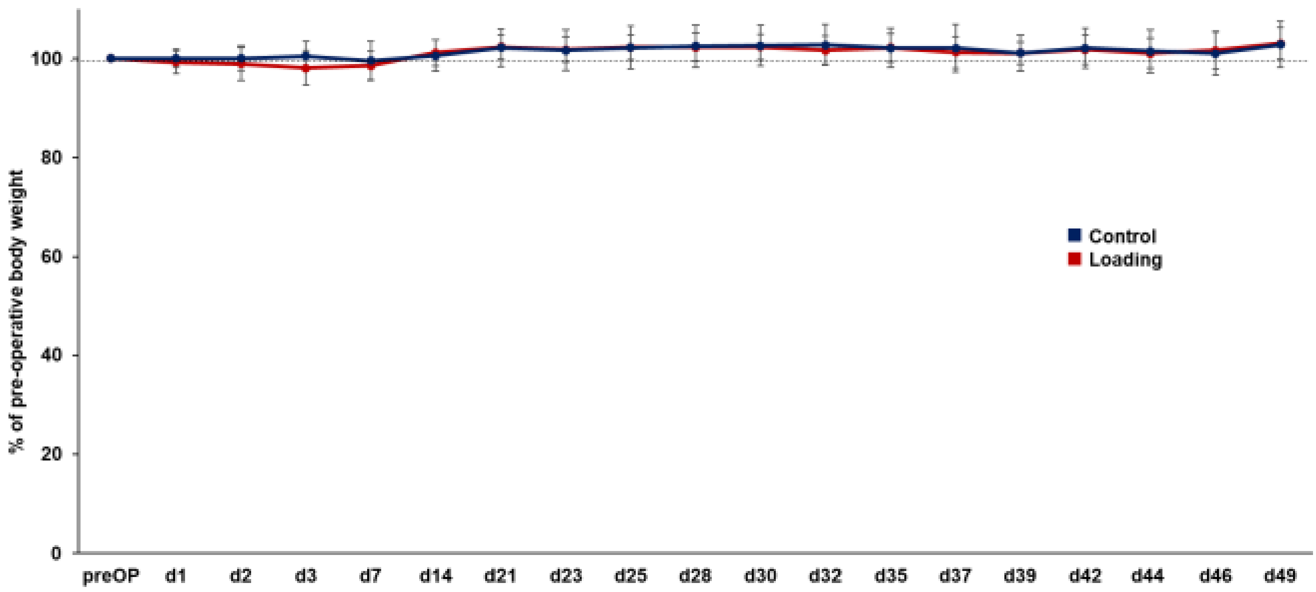


**Supplementary Fig. S2.** *In vivo* monitoring of body weight of the mice from the control group (n=10) and the loading group (n=10) measured pre-operatively (preOP), on postoperative days 1-3, weekly from day 7 to day 14 and 3x/week during the loading period (day 21 - day 49). The postoperative values were related to the preoperative data.

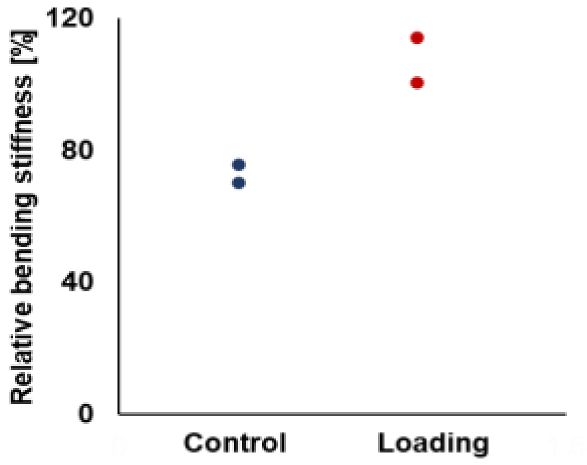


**Supplementary Fig. S3.** Relative bending stiffness (healed femur vs. contralateral femur) was assessed via *ex vivo* 3-point-bending testing for two animals from the loading and the control group.

**Supplementary Table S1.** Study design (female 20 week-old C57BL/6J mice)

| Group | Femur defect saw size: 0.66mm | *In vivo* micro-CT measurements | Registration of micro-CT scans # | Loading | Histology |
|---|---|---|---|---|---|
| Control | n=10 | d0, week 1-7 (n=10) | week 1-7 (n=10) | 0N, 3/week, week 3-7 (n=10) | week 7 (n=1) |
| Loading | n=10 | d0, week 1-7 (n=10) | week 1-7 (n=10) | 8-16N, 10 Hz, 3/week, week 3-7 (n=10) | week 7 (n=1) |

### micro-CT scan taken at timepoint x registered to micro-CT scan taken at timepoint x-1
